## Supplementary material for "Removal of reinforcement improves instrumental performance in humans by decreasing a general action bias rather than unmasking learnt associations": S1Appendix

### Contents

1. RESULTS SEPARATELY FOR THE TWO EXPERIMENTS 2

1.1. Experiment 1 2

1.2. Experiment 2 7

2. BEHAVIOURAL ANALYSIS OF THE POOLED DATA SET 12

3. ANALYSIS OF MODELLING RESULTS 15

3.1. Model comparison with a set of fixed learning rates 15

3.2. Parameter recovery for the bias model 16

3.3. Model validation 17

3.3.1. Individual parameters 17

3.3.2. Types of forgetting 19

3.3.3. Modelling the behaviour in probe trials 20

4. CONTROL ANALYSIS FOR REACTION TIMES 22

### 1. Results separately for the two experiments

We acquired two independent data sets (each *N* = 30), which we pooled for the main analyses. Here, we present the results separately for Experiment 1 (Fig 1-3 and Table 1-4) and Experiment 2 (Fig 4-6 and Table 5-8).

#### 1.1. Experiment 1

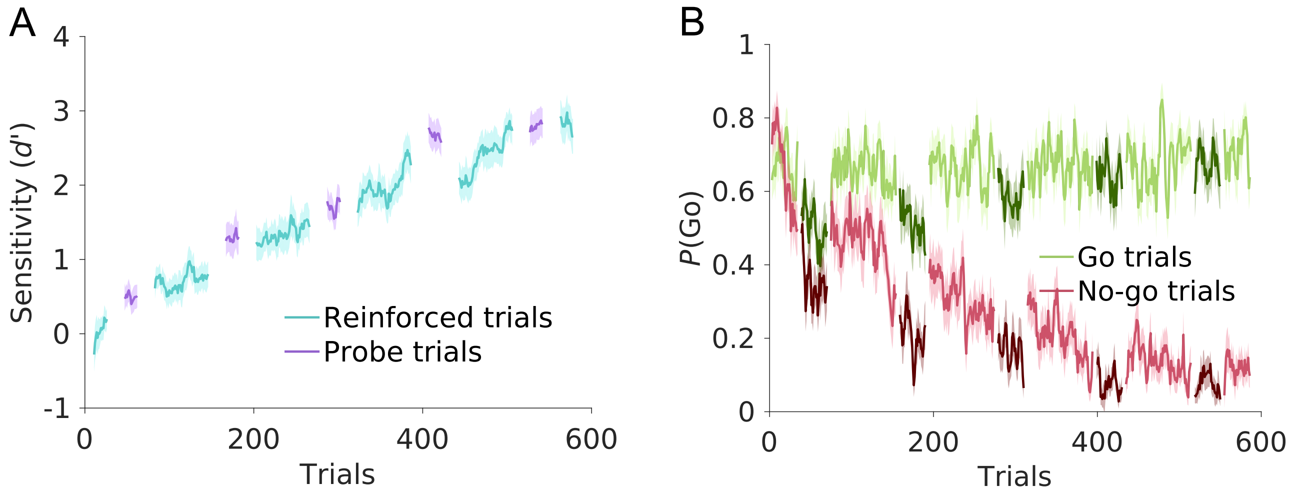


**Fig 1. Average learning performance for Experiment 1 (*N* = 30).**
(A) Sensitivity index *d'*, separately for reinforced (cyan) and probe trials (purple). Solid lines represent mean performance, shaded areas SEM across participants. (B) Time course of go-response probabilities, *P*(Go), for go trials (green) and no-go trials (red). Darker shades of green and red illustrate probe trials. Solid lines represent mean, shaded areas SEM across participants.

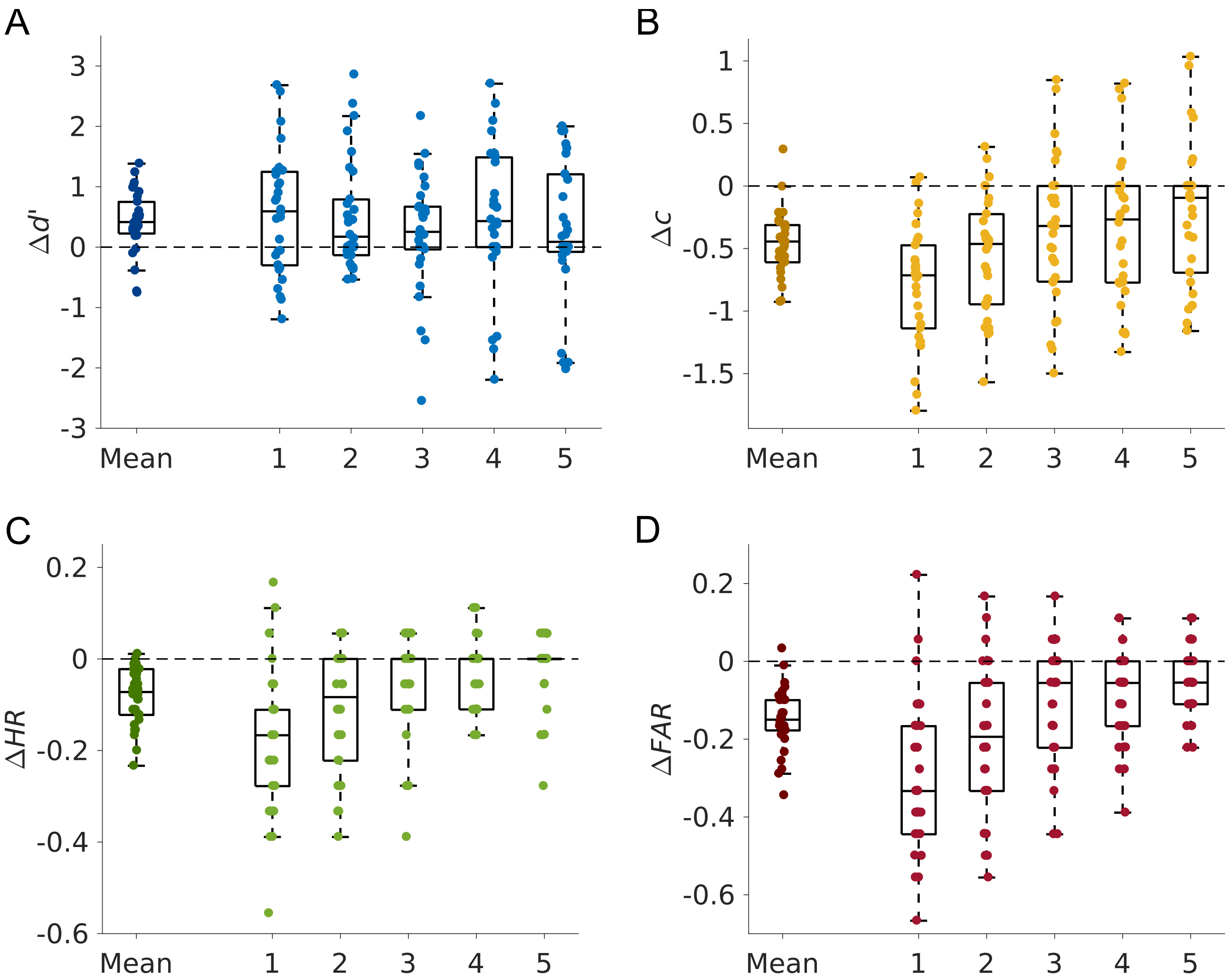


**Fig 2. Behavioural results, expressed as difference between probe trials and preceding reinforced trials for Experiment 1 (*N* = 30).**

Results are shown both for the mean across all five probe blocks (left) and separately for each probe block. (A**)** The sensitivity index *d'* increased in probe compared to reinforced trials. (B) The negative bias criterion *c*, decreased in probe blocks, indicating a reduced propensity to act on probe trials. (C), (D) Both hit rate (*HR*, C) and false alarm rate (*FAR*, D) decreased in probe blocks.

**Table 1. Difference between probe and pre-probe trials for Experiment 1.**
Statistical comparisons were performed using Student's *t*-test.

| | Mean | *SEM* | $t_{29}$ | *p* | Cohen's *d* |
|---|---|---|---|---|---|
| *d'* | 0.42 | 0.50 | 4.57 | <.001 | 0.83 |
| *c* | -0.46 | 0.25 | -9.84 | <.001 | -1.807 |
| *HR* | -0.08 | 0.06 | -7.11 | <.001 | -1.309 |
| *FAR* | -0.15 | 0.08 | -10.16 | <.001 | -1.86 |

**Table 2. Effect of time on the difference between probe and pre-probe trials for Experiment 1.**
Statistical comparisons were performed using repeated measures ANOVA with Greenhouse-Geisser correction, where appropriate.

| | $F(4, 29)$ | *p* | $\eta^2$ |
|---|---|---|---|
| *d'* | 0.58 | .675 | 0.02 |
| *c* | 5.47 | <.001 | 0.16 |
| *HR* | 10.15 | <.001 | 0.26 |
| *FAR* | 14.09 | <.001 | 0.33 |

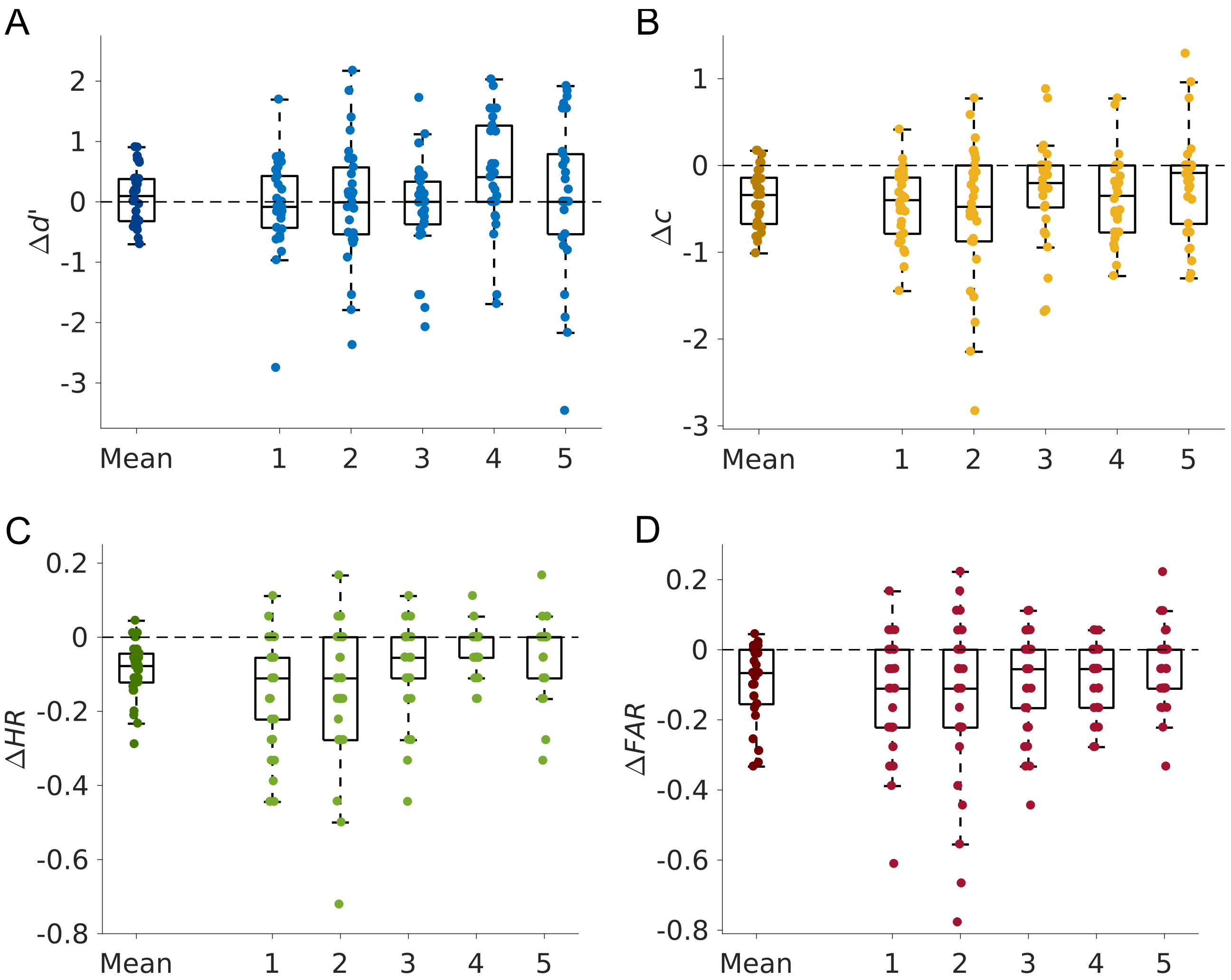


**Fig 3. Behavioural results, expressed as difference between probe trials and subsequent reinforced trials for Experiment 1 (*N* = 30).**

Results are shown both for the mean across all five probe blocks (left) and separately for each probe block. (A) The sensitivity index *d'* was not significantly different in probe compared to reinforced trials. (B) The negative bias criterion *c*, decreased in probe blocks, indicating a reduced propensity to act on probe trials. (C), (D) Both hit rate (*HR*, C) and false alarm rate (*FAR*, D) decreased in probe blocks.

**Table 3. Difference between probe and post-probe trials for Experiment 1.**
Statistical comparisons were performed using Student's *t*-test.

| | Mean | *SEM* | $t_{29}$ | *p* | Cohen's *d* |
|---|---|---|---|---|---|
| *d'* | 0.08 | 0.46 | 0.90 | .375 | 0.17 |
| *c* | -0.38 | 0.34 | -6.05 | <.001 | -1.11 |
| *HR* | -0.09 | 0.08 | -6.41 | <.001 | -1.17 |
| *FAR* | -0.10 | 0.11 | -4.89 | <.001 | -0.89 |

**Table 4. Effect of time on the difference between probe and post-probe trials for Experiment 1.**
Statistical comparisons were performed using repeated measures ANOVA with Greenhouse-Geisser correction, where appropriate.

| | $F(4, 29)$ | *p* | $\eta^2$ |
|---|---|---|---|
| *d'* | 1.82 | .130 | 0.06 |
| *c* | 1.76 | .141 | 0.06 |
| *HR* | 5.83 | .001 | 0.17 |
| *FAR* | 3.32 | .028 | 0.10 |

### 1.2. Experiment 2

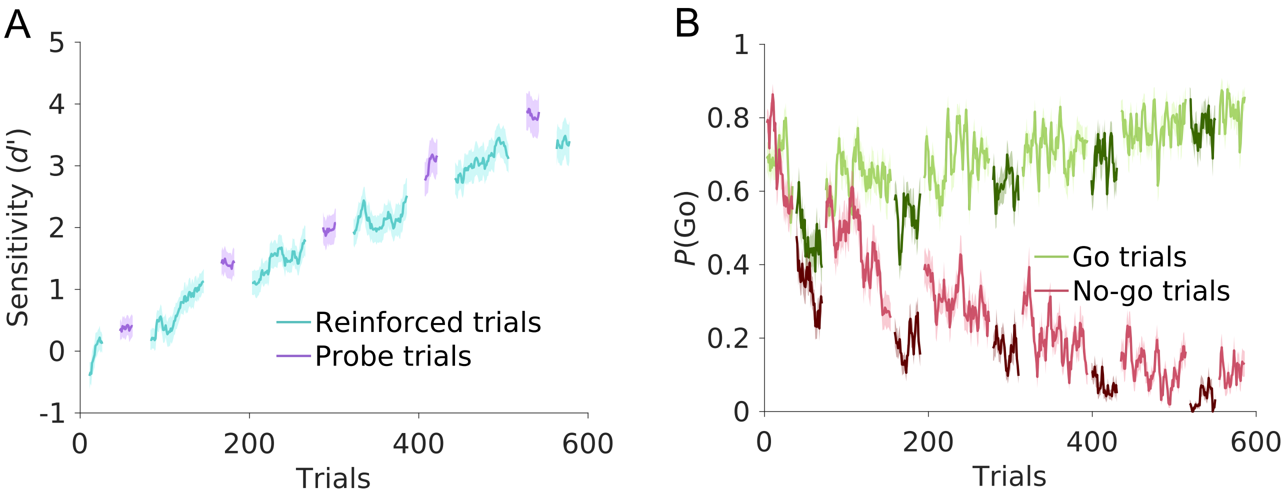


**Fig 4. Average learning performance for Experiment 2 (*N* = 30).**
(A) Sensitivity index *d'*, separately for reinforced (cyan) and probe trials (purple). Solid lines represent mean performance, shaded areas SEM across participants. (B) Time course of go-response probabilities, *P*(Go), for go trials (green) and no-go trials (red). Darker shades of green and red illustrate probe trials. Solid lines represent mean, shaded areas SEM across participants.

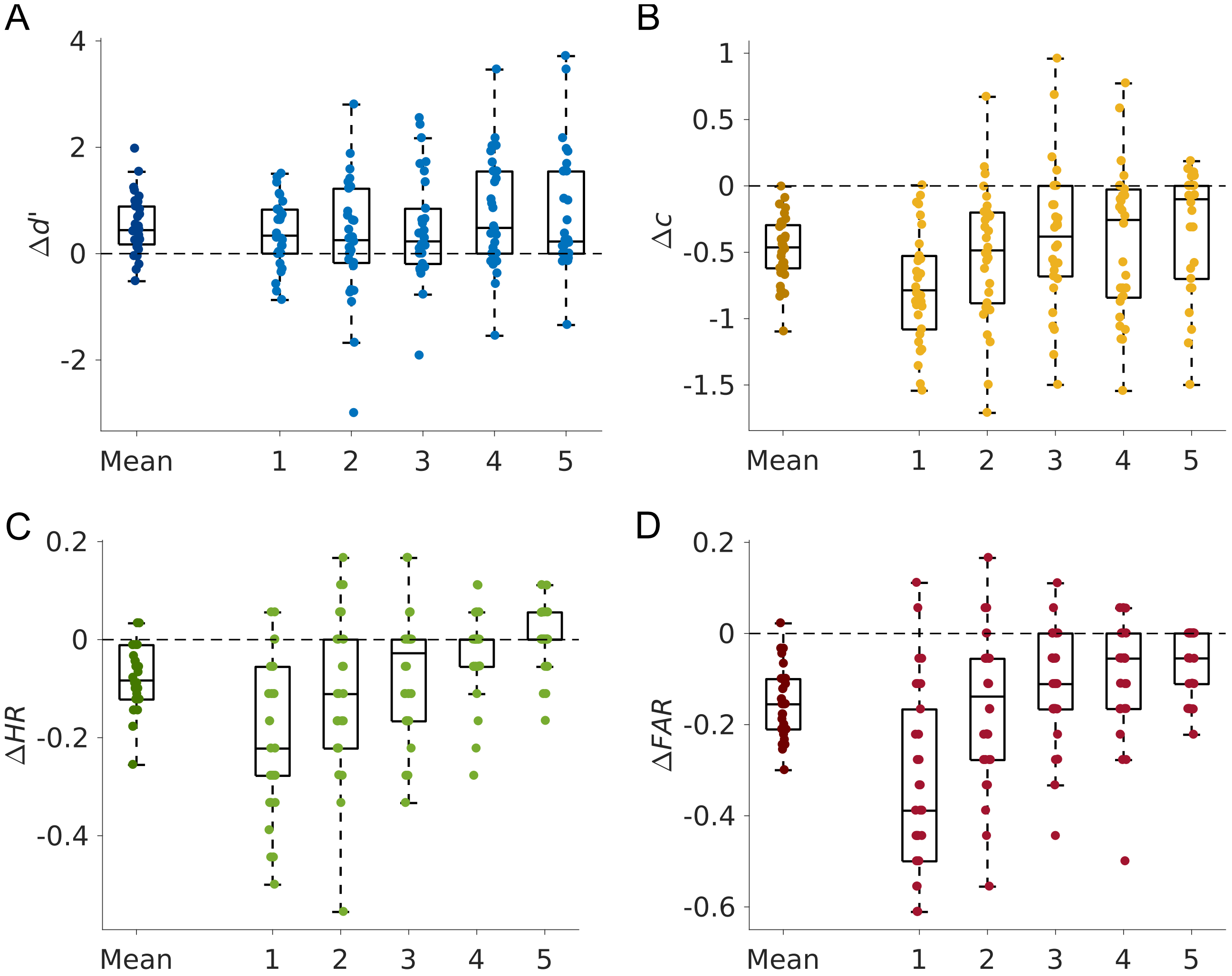


**Fig 5. Behavioural results, expressed as difference between probe trials and preceding reinforced trials for Experiment 2 (*N* = 30).**

Results are shown both for the mean across all five probe blocks (left) and separately for each probe block. (A) The sensitivity index *d'* increased in probe compared to reinforced trials. (B) The negative bias criterion *c*, decreased in probe blocks, indicating a reduced propensity to act on probe trials. (C), (D) Both hit rate (*HR*, C) and false alarm rate (*FAR*, D) decreased in probe blocks.

**Table 5. Difference between probe and pre-probe trials for Experiment 2.**
Statistical comparisons were performed using Student's *t*-test.

| | Mean | *SEM* | $t_{29}$ | *p* | Cohen's *d* |
|---|---|---|---|---|---|
| *d'* | 0.52 | 0.55 | 5.19 | <.001 | 0.95 |
| *c* | -0.48 | 0.25 | -10.61 | <.001 | -1.94 |
| *HR* | -0.08 | 0.07 | -6.48 | <.001 | -1.18 |
| *FAR* | -0.15 | 0.08 | -11.08 | <.001 | -2.02 |

**Table 6. Effect of time on the difference between probe and pre-probe trials for Experiment 2.**
Statistical comparisons were performed using repeated measures ANOVA with Greenhouse-Geisser correction, where appropriate.

| | $F(4, 29)$ | *p* | $\eta^2$ |
|---|---|---|---|
| *d'* | 1.44 | .227 | 0.05 |
| *c* | 3.72 | .007 | 0.11 |
| *HR* | 13.91 | <.001 | 0.32 |
| *FAR* | 18.81 | <.001 | 0.39 |

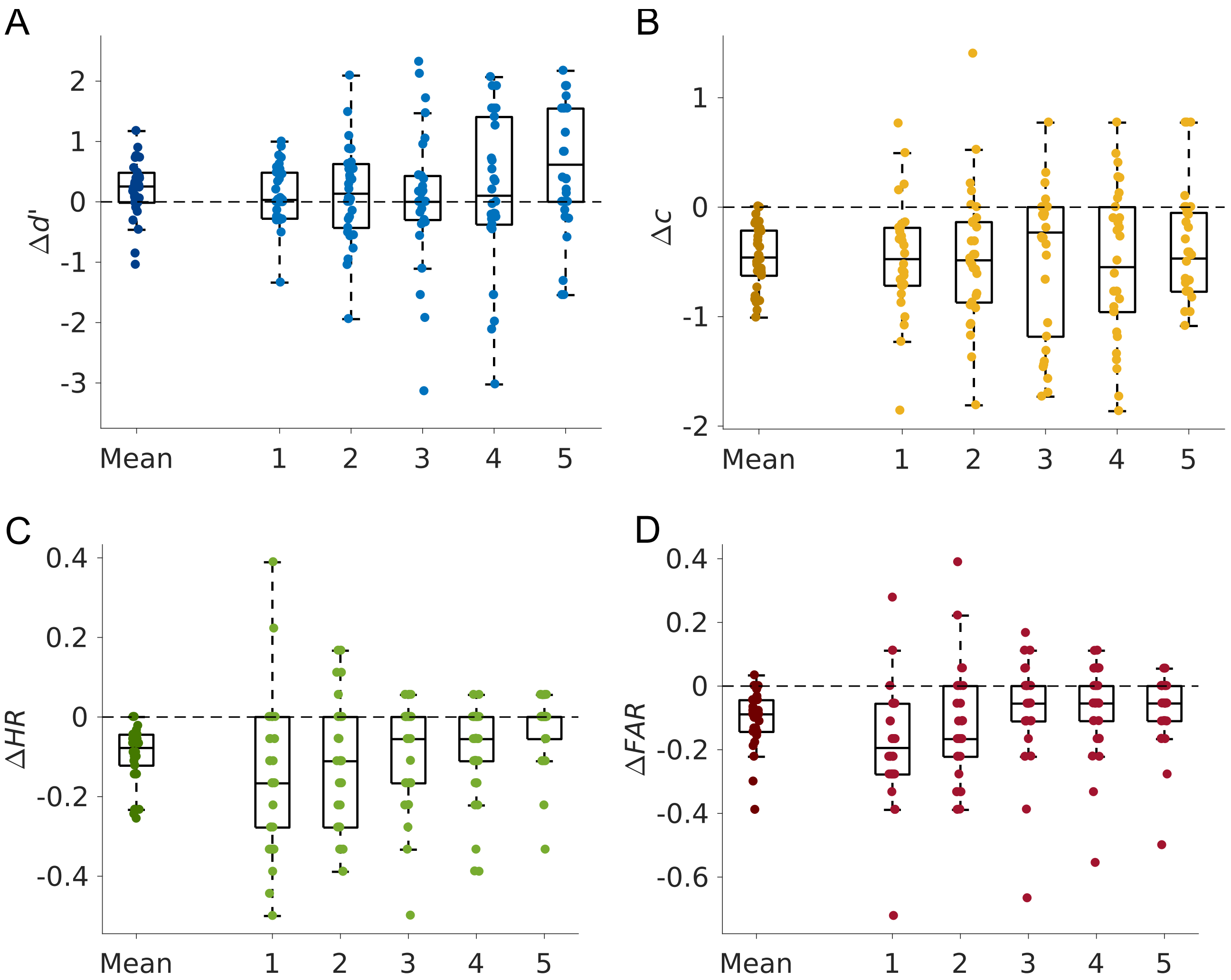


**Fig 6. Behavioural results, expressed as difference between probe trials and subsequent reinforced trials for Experiment 2 (*N* = 30).**

Results are shown both for the mean across all five probe blocks (left) and separately for each probe block. (A) The sensitivity index *d’* increased in probe compared to reinforced trials. (B) The negative bias criterion *c*, decreased in probe blocks, indicating a reduced propensity to act on probe trials. (C), (D) Both hit rate (*HR*, C) and false alarm rate (*FAR*, D) decreased in probe blocks.

**Table 7. Difference between probe and post-probe trials for Experiment 2.**
Statistical comparisons were performed using Student's *t*-test.

| | Mean | *SEM* | $t_{29}$ | *p* | Cohen's *d* |
|---|---|---|---|---|---|
| *d'* | 0.22 | 0.48 | 2.47 | .020 | 0.45 |
| *c* | -0.46 | 0.29 | -8.64 | <.001 | -1.58 |
| *HR* | -0.09 | 0.07 | -7.40 | <.001 | -1.35 |
| *FAR* | -0.11 | 0.09 | -6.47 | <.001 | -1.18 |

**Table 8. Effect of time on the difference between probe and post-probe trials for Experiment 2.**
Statistical comparisons were performed using repeated measures ANOVA with Greenhouse-Geisser correction, where appropriate.

| | $F(4, 29)$ | *p* | $\eta^2$ |
|---|---|---|---|
| *d'* | 1.70 | .155 | 0.06 |
| *c* | 0.24 | .915 | 0.01 |
| *HR* | 3.13 | .040 | 0.10 |
| *FAR* | 3.50 | .018 | 0.11 |

The results for both experiments are very similar. The comparison of pre-probe and probe trials yield identical results for both experiments: *d'*, bias criterion, hit and false alarm rate change significantly and there is a significant effect of time for these parameters except for *d'*. The results of the comparison of post-probe and probe trials are similar for both experiments, with the exception that *d'* decreased significantly in post-probe trials in Experiment 2, but not in Experiment 1.

### 2. Behavioural analysis of the pooled data set

Here, we report the post hoc tests for the difference in the sensitivity *d'* between probe and pre-probe trials (Table 9). Additionally, we analysed the difference between probe and post-probe trials of the pooled data (Fig 7 and Table 10).

**Table 9. Post hoc tests for the difference in *d'* of probe and pre-probe trials.**

| Block | $t_{59}$ | *p* | Cohen's *d* |
|---|---|---|---|
| 1 | 4.44 | <.001 | 0.57 |
| 2 | 3.03 | .004 | 0.39 |
| 3 | 2.74 | .008 | 0.35 |
| 4 | 4.35 | <.001 | 0.56 |
| 5 | 3.43 | .001 | 0.44 |

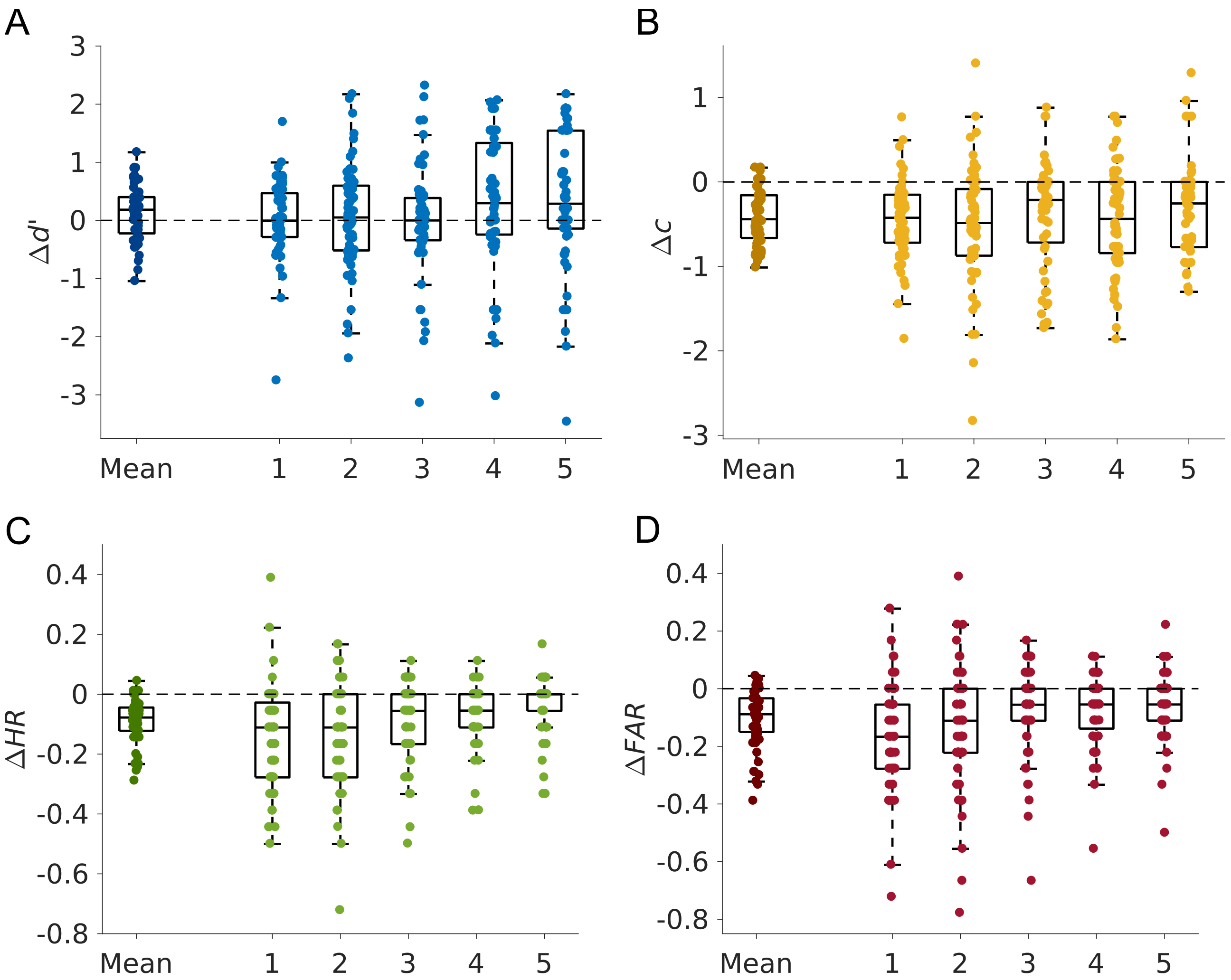


**Fig 7. Behavioural results, expressed as difference between probe trials and subsequent reinforced trials for all participants (*N* = 60).**

Results are shown both for the mean across all five probe blocks (left) and separately for each probe block. (A) The sensitivity index *d'* increased in probe compared to reinforced trials. (B) The negative bias criterion *c*, decreased in probe blocks, indicating a reduced propensity to act on probe trials. (C), (D) Both hit rate (*HR*, C) and false alarm rate (*FAR*, D) decreased in probe blocks.

**Table 10. Effect of time on the difference between probe and post-probe trials for all participants.**
Statistical comparisons were performed using repeated measures ANOVA with Greenhouse-Geisser correction, where appropriate.

| | $F(4, 59)$ | $p$ | $\eta^2$ |
|---|---|---|---|
| *d'* | 1.99 | .097 | 0.03 |
| *c* | 1.22 | .304 | 0.02 |
| *HR* | 7.66 | .001 | 0.12 |
| *FAR* | 6.21 | .001 | 0.10 |

### 3. Analysis of modelling results

We compared all models with five additional learning rates evenly log-spaced between 0.01 and 0.2 to verify that that the results are not dependent on the learning rate $\alpha = 0.06$, which we chose for the main analysis (Table 11 and Fig 8). The results of the recovery for all free parameters of the bias model are shown in this part (Fig 9 and Table 12).

#### 3.1. Model comparison with a set of fixed learning rates

**Table 11. Comparisons of baseline, temperature, bias and full model with varying learning rates $\boldsymbol{\alpha}$.** Parameters were fit for all participants (*N* = 60), and *BIC*s (mean ± SEM) were calculated for model comparison. Learning rate $\alpha$ = 0.06 which was used for main analyses is bold.

| $\alpha$ | Baseline model | Temperature model | Bias model | Full model |
|---|---|---|---|---|
| 0.01 | 512.47 ± 134.65 | 515.60 ± 134.34 | 504.87 ± 129.09 | 509.71 ± 129.17 |
| 0.018 | 518.10 ± 135.23 | 521.07 ± 134.82 | 510.76 ± 129.67 | 515.45 ± 129.71 |
| 0.033 | 526.96 ± 135.96 | 529.89 ± 135.65 | 520.08 ± 130.47 | 524.67 ± 130.53 |
| **0.06** | **539.42 ± 137.39** | **542.22 ± 137.25** | **533.25 ± 131.99** | **537.56 ± 132.10** |
| 0.11 | 555.32 ± 138.50 | 557.98 ± 138.56 | 550.02 ± 133.19 | 554.08 ± 133.32 |
| 0.2 | 573.18 ± 138.01 | 575.43 ± 137.39 | 568.69 ± 133.17 | 572.56 ± 132.85 |

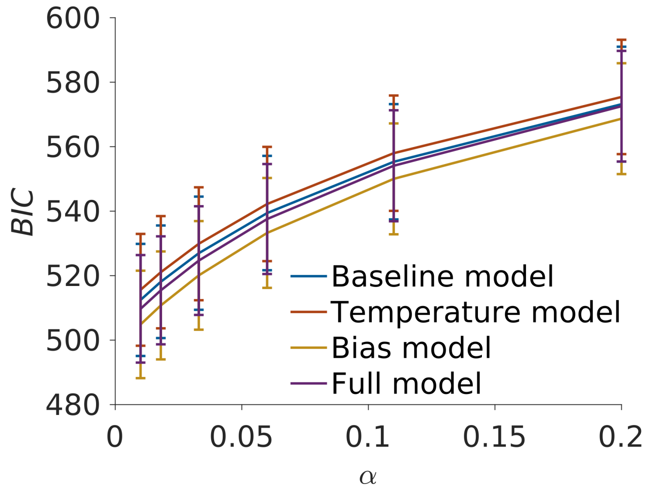


**Fig 8. Model comparison with different learning rates.**
Comparison of mean *BIC*s for six different learning rates, error bars represent SEM. For all learning rates, the bias model provided the best fit. The full model fitted second best, followed by the baseline model. The temperature model performed the worst.

#### 3.2. Parameter recovery for the bias model

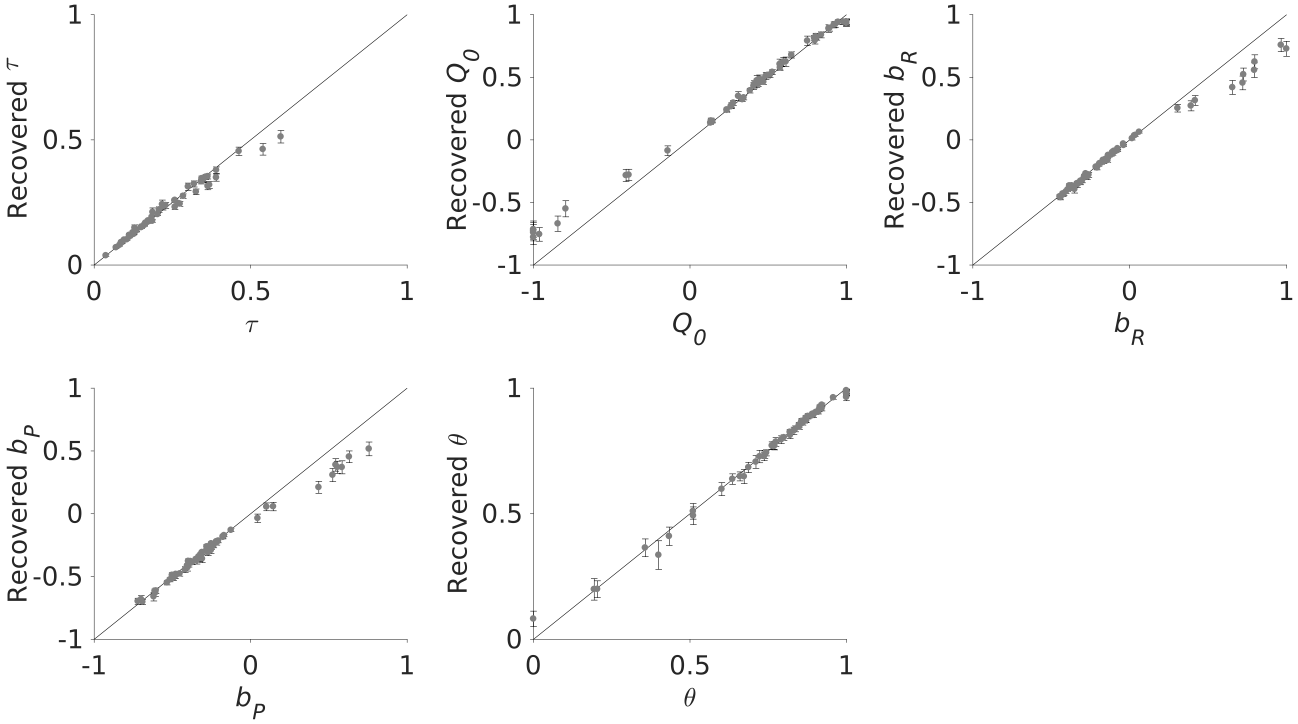


**Fig 9. Parameter recovery for all free parameters of the bias model.**
Parameters fitted to participants' behaviour are plotted against the recovered parameters. Error bars represent 95% Cousineau-Morey confidence intervals.

**Table 12. Correlation coefficients for the recovery of each parameter.**

| Free Parameter | $\rho$ | $p$ |
|---|---|---|
| $\tau$ | 0.991 | <.001 |
| $Q_0$ | 0.998 | <.001 |
| $b_R$ | 0.994 | <.001 |
| $b_P$ | 0.995 | <.001 |
| $\theta$ | 0.997 | <.001 |

#### 3.3. Model validation

Because of the artefacts of the SDT analysis, we did not have quantitative measures for the model validation. Therefore, we plotted the go-response probabilities based on model simulations and inspected visually whether the goodness of model fit supports the *BIC* outcomes. First, we validated whether the individual parameters of the baseline model are necessary to describe the general behaviour (Fig 10). Second, we compared different types of forgetting (Fig 11). Third, we validated whether the temperature or bias model could reproduce the behavioural change in probe trials (Fig 12).

##### 3.3.1. Individual parameters

First, we ignored the probe trials and checked whether all parameters in the baseline model are needed to describe the empirical behaviour: Participants' go-response probabilities for both go and no-go trials started high with the probability for go trials staying high and the probability for no-go trials decreasing over time.

We started with a model containing two free parameters: a softmax temperature and a general bias. This simple model was not suitable to describe the empirical data as the go-response probabilities start relatively low and the probabilities for go and no-go trials increased and decreased over time, respectively (*BIC* = 608.47 ± 118.63, mean ± SEM, Fig 10A). Adding an initial *Q*-value improved the fit compared to the simplest model, but was still not able to reproduce the participants' behaviour (*BIC* = 595.62 ± 119.28, Fig 10B). The same applies to a model containing of a softmax temperature, a general bias and a decay parameter (*BIC* = 591.77 ± 123.25, Fig 10C). In this model, we assumed a decay of learnt values towards zero when no go-response is performed. For the baseline model, we combined a softmax temperature, a general bias, an initial *Q*-value and a decay parameter and this model is able to reproduce the behaviour described above (*BIC* = 539.42 ± 137.39, Fig 10D).

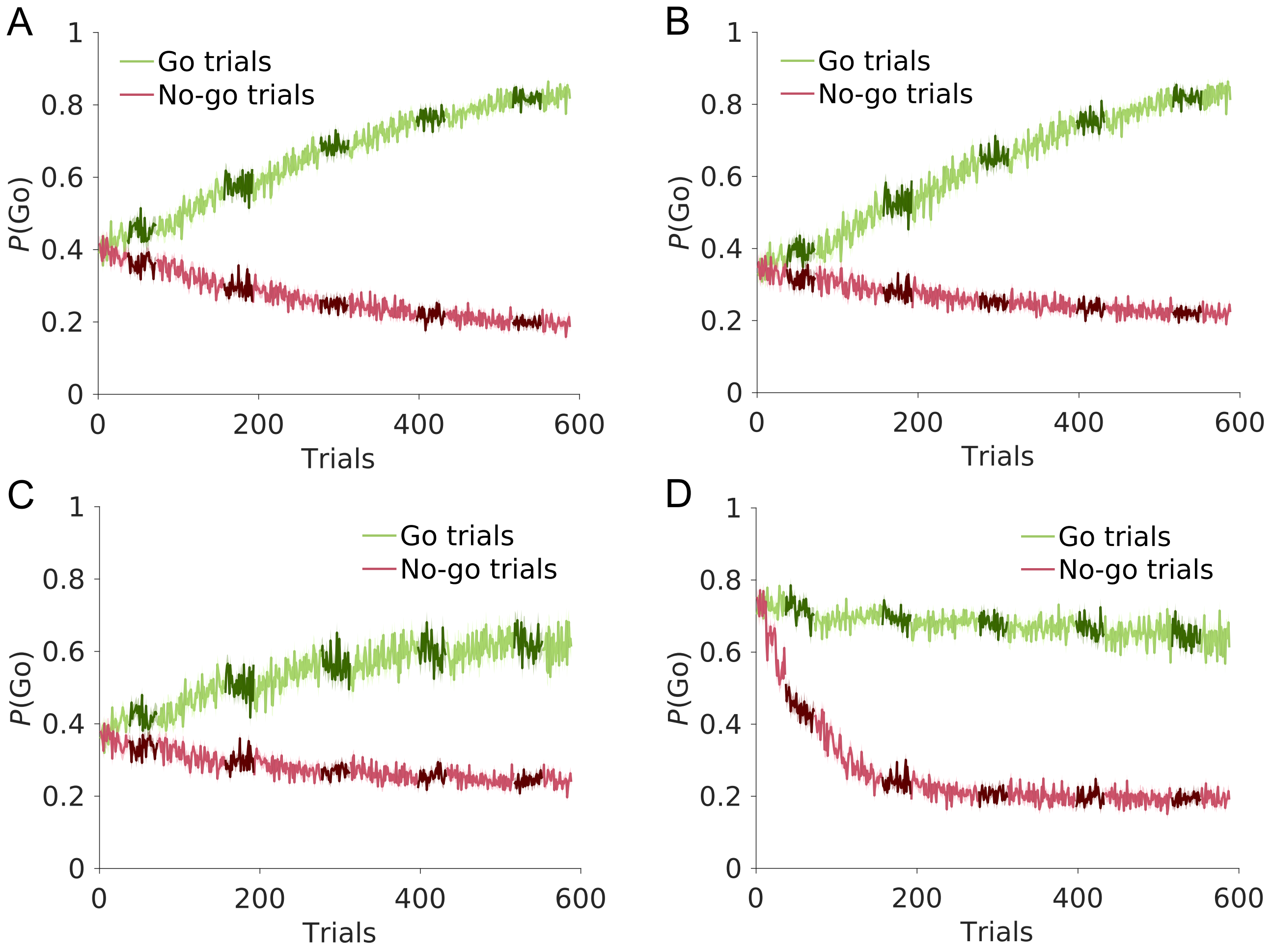


**Fig 10. Set up of the baseline model.**
Time course of simulated go-response probabilities, *P*(Go), for go trials (green) and no-go trials (red). Darker shades of green and red illustrate probe trials. Solid lines represent mean, shaded areas SEM across simulations. Simulations are based on (A) a softmax temperature and a general bias, (B) a softmax temperature, a general bias and an initial *Q*-value, (C) a softmax temperature, a general bias and a decay parameter and (D) the complete baseline model (softmax temperature, general bias, initial *Q*-value, decay parameter).

#### 3.3.2. Types of forgetting

There are several ways to implement a decay of option values due to forgetting. We implemented two different ways of forgetting and compared it to our baseline model. First, we set up a model in which values decay towards the initial *Q*-value. The model worsened again and due to a low initial *Q*-value, the go-response probabilities start low with the probabilities for no-go trials staying low and the probabilities for go trials increasing over time, which is not in line with the participants' behaviour (*BIC* = 547.74 ± 116.46, Fig 11A).

Another approach for the decay parameter is to implement forgetting when no feedback for the go-response is received (instead of forgetting after no-go-responses). Again, this model performed worse compared to the baseline model and could not capture the participants' behaviour (*BIC* = 600.70 ± 117.24, Fig 11B).

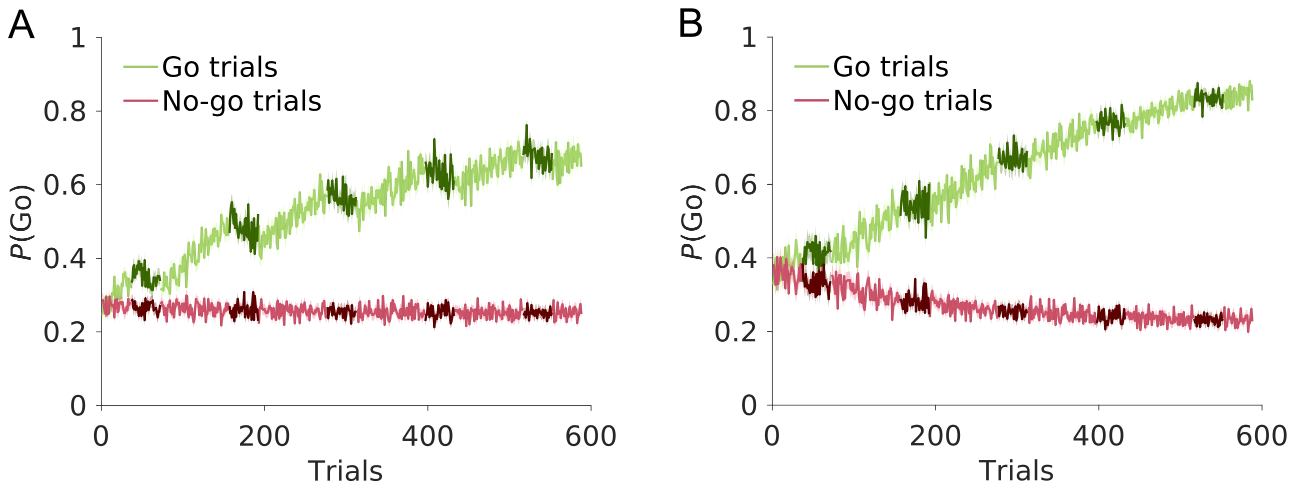


**Fig 11. Variations of the baseline model.**
Time course of simulated go-response probabilities, *P*(Go), for go trials (green) and no-go trials (red). Darker shades of green and red illustrate probe trials. Solid lines represent mean, shaded areas SEM across simulations. Simulations are based on (A) a model with values decaying towards the initial *Q*-value instead of zero and (B) a model with values decaying when no feedback is given instead of no response decay.

#### 3.3.3. Modelling the behaviour in probe trials

In probe trials, participants' go-response probabilities for both go and no-go trials decreased. Based on the baseline model, we now implemented two models differentiating between reinforced and probe trials; In the temperature model fitted a softmax temperature separately for each trial type. It performed worse than the baseline model and the decrease in go-response probabilities for both go and no-go trials is not comparable to the observed behaviour (*BIC* = 542.22 ± 137.25, Fig 12A). The bias model, which fitted a general bias separately for each trial type, performed better than the baseline model and looks comparable to the participants' behaviour (*BIC* = 533.25 ± 131.99, Fig 12B).

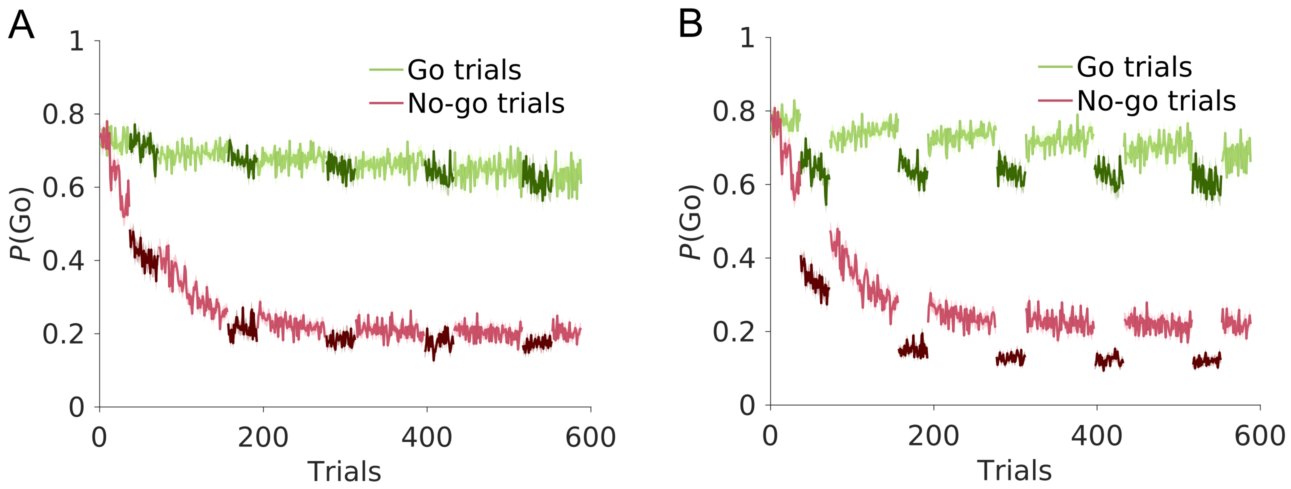


**Fig 12. Simulations of changed behaviour in probe trials.**

Time course of simulated go-response probabilities, *P*(Go), for go trials (green) and no-go trials (red). Darker shades of green and red illustrate probe trials. Solid lines represent mean, shaded areas SEM across simulations. Simulations are based on (A) the temperature model and (B) the bias model.

#### 4. Control analysis for reaction times

To ensure that the response window was long enough for participants to administer a go-response, we plotted the distribution of participants' reaction times (RTs) in all trials.

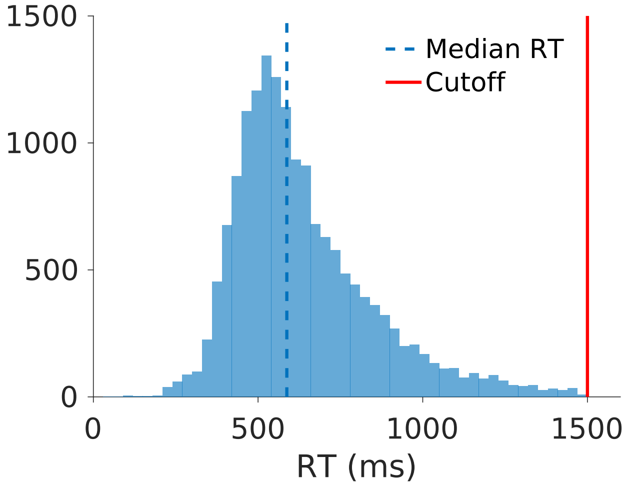


**Fig 13. Distribution of participants' RTs.**
The cutoff of 1500 ms relative to stimulus onset marks the end of the response window.
